## Supplemental data for "The MODY-associated *KCNK16* L114P mutation increases islet glucagon secretion and limits insulin secretion resulting in transient neonatal diabetes and glucose dyshomeostasis in adults"

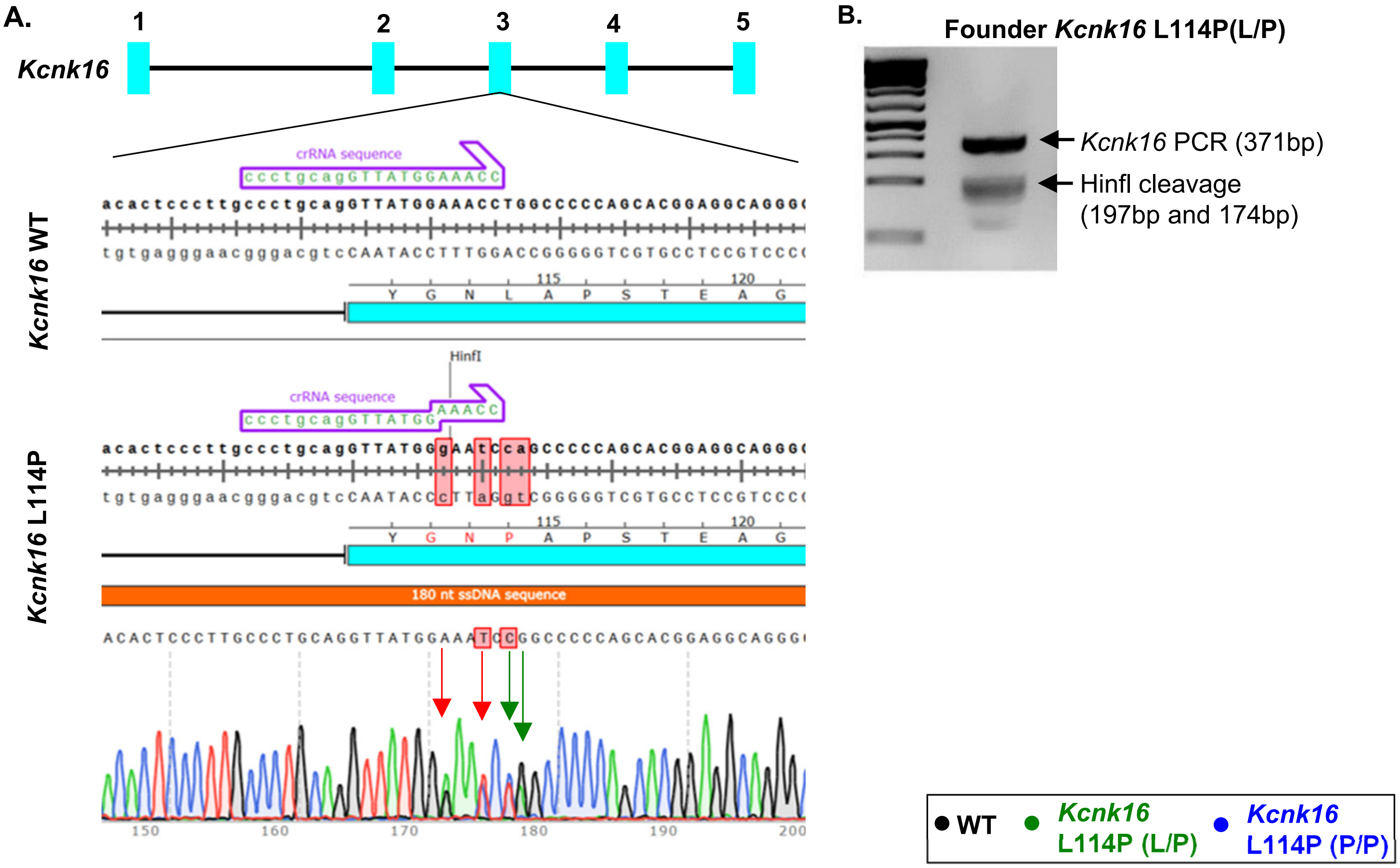


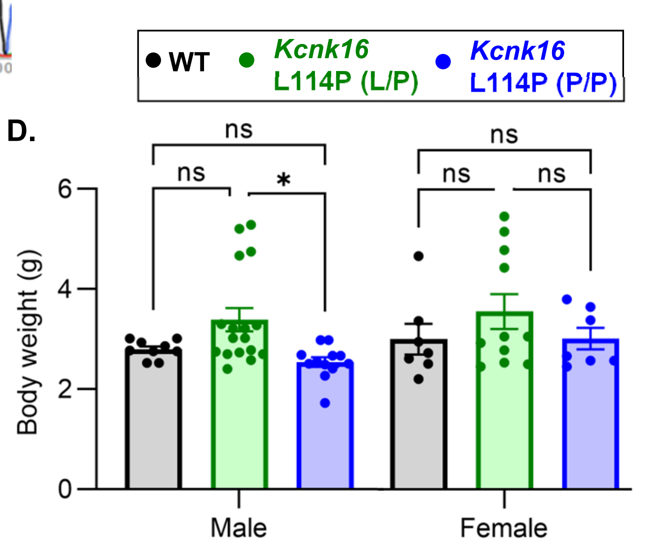


C.

| Genotype | Expected | Observed |
|---|---|---|
| *Kcnk16* WT | 25% | 27% |
| *Kcnk16* L114P (L/P) | 50% | 53% |
| *Kcnk16* L114P (P/P) | 25% | 6.3% |

$\chi^2$ = 6.648, *P* = 0.036

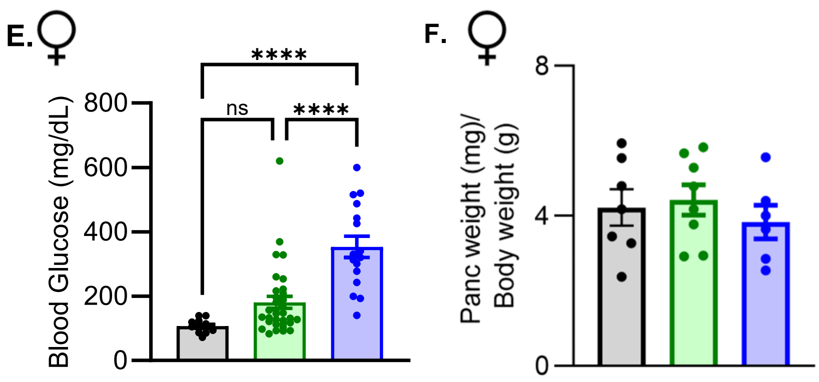

**Supplemental figure 1. Generation of *Kcnk16* L114P model and assessment of neonatal glucose homeostasis and lethality. A.** Targeted region of *Kcnk16* exon 3 using CRISPR/Cas9 leading to the introduction of HinfI restriction enzyme site (red arrows) and CTG to CCA mutation in codon 337 corresponding to p.TALK-1 L114P (green arrows). **B.** PCR confirmation of the male founder *Kcnk16* L114P (L/P) mouse using HinfI restriction digestion. **C.** $\chi^2$ analysis of the F1 progeny from B6; CD-1 *Kcnk16* L114P (L/P) crosses. **D.** Body weight measurements of male (left) and female (right) wildtype (WT; black), heterozygous *Kcnk16* L114P (L/P; green), and homozygous *Kcnk16* L114P (P/P; blue) mice on P4. **E.** Blood glucose measurements of female mice on P4. **F.** Pancreas weight/ body weight measurements of P4 female mice. Data are presented as mean±SEM. Data were analyzed using student's t-test or one-way ANOVA.

A.

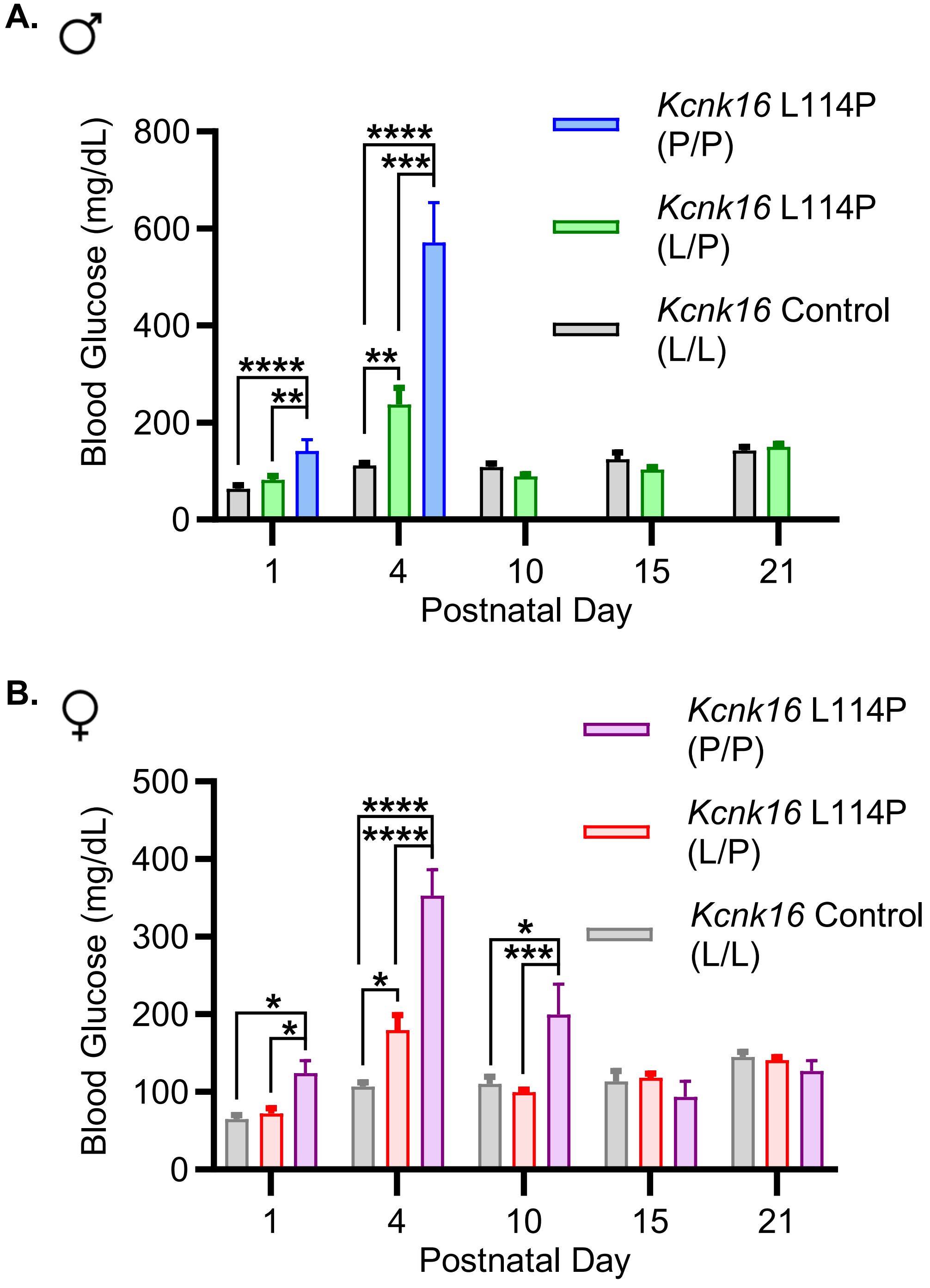
♂
Blood Glucose (mg/dL)
800
600
400
200
0
****
***
**
****
**
1
4
10
15
21
Postnatal Day
Kcnk16 L114P (P/P)
Kcnk16 L114P (L/P)
Kcnk16 Control (L/L)
B.
♀
500
400
300
200
100
0
****
****
*
*
***
*
*
Kcnk16 L114P (P/P)
Kcnk16 L114P (L/P)
Kcnk16 Control (L/L)

**Supplemental figure 2.** ***Kcnk16*** **L114P male and female mice show transient neonatal hyperglycemia. A.** Average blood glucose measurements of B6;CD-1 male mice at P1, P4, P10, P15, and P21 including: WT (black; P1 N=9, P4 N=27, P10 N=9, P15 N=9, P21 N=8), *Kcnk16* L114P (L/P; green; P1 n=15, P4 N=28, P10 N=24, P15 N=14, P21 N=14), and *Kcnk16* L114P (P/P; green; P1 n=5, P4 N=10, no P/P male mice survived past P4). **B.** Average blood glucose measurements of B6;CD-1 female mice at P1, P4, P10, P15, and P21 including: WT (black; P1 N=8, P4 N=13, P10 N=8, P15 N=8, P21 N=8), *Kcnk16* L114P (L/P; green; P1 n=24, P4 N=33, P10 N=24, P15 N=24, P21 N=24), and *Kcnk16* L114P (P/P; green; P1 n=10, P4 N=16, P10 N=5, P15 N=4, P21 N=4). Data are presented as mean±SEM. Data were analyzed using student's t-test; *P<0.05, **P<0.01, ***P<0.001, ***P<0.0001.

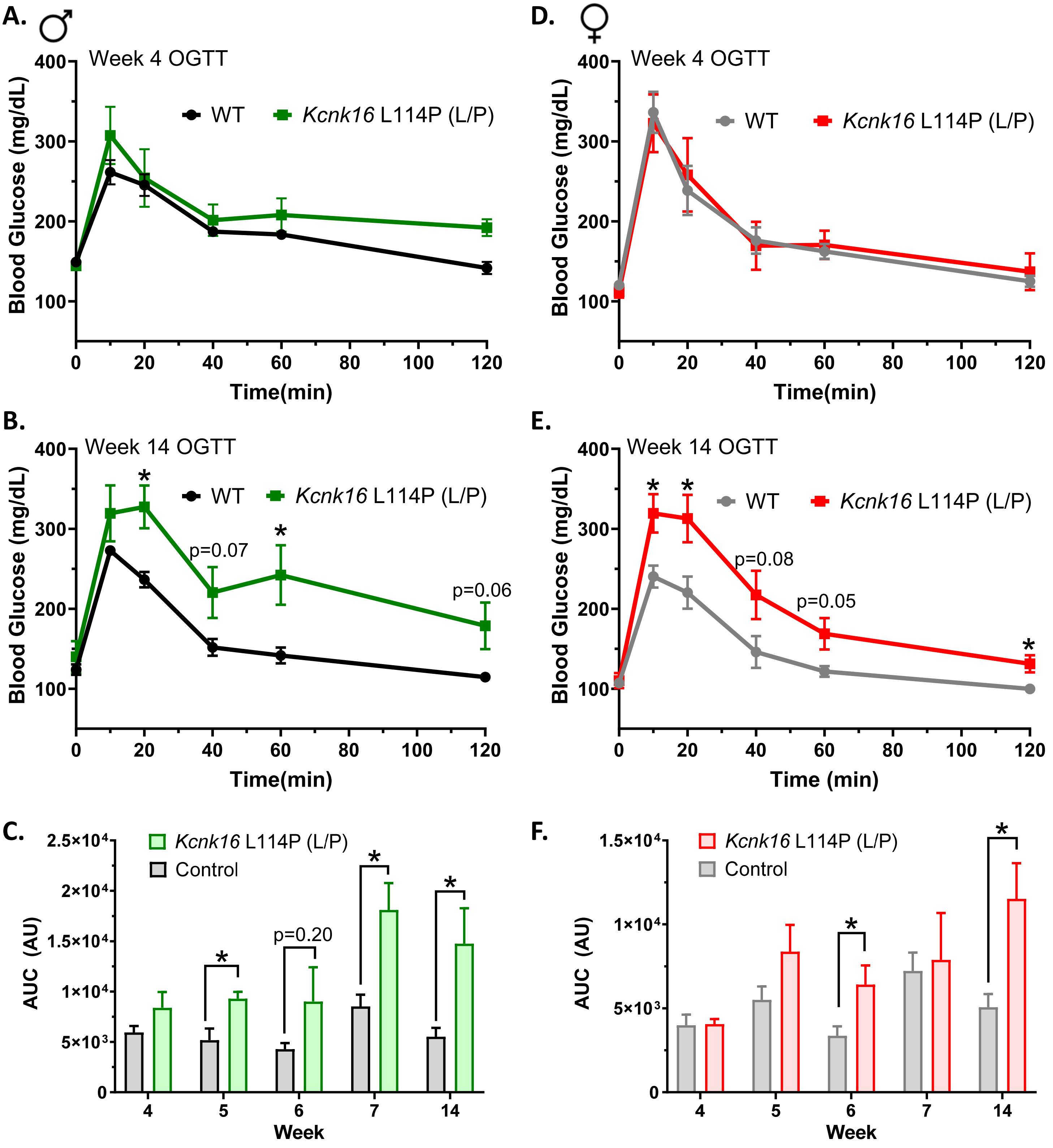

A. ♂
Week 4 OGTT
WT
Kcnk16 L114P (L/P)
Blood Glucose (mg/dL)
Time(min)
B.
Week 14 OGTT
p=0.07
p=0.06
C.
Kcnk16 L114P (L/P)
Control
AUC (AU)
p=0.20
Week
D. ♀
Week 4 OGTT
E.
Week 14 OGTT
p=0.08
p=0.05
Time (min)
F.

**Supplemental figure 3. Oral glucose tolerance is impaired in *Kcnk16* L114P (L/P) mice in the B6;CD-1 background.** Oral glucose tolerance test (OGTT) performed in 4 (**A**) and 14-week-old (**B**) male mice following a 4-hour fast in response to oral gavage of 2g/kg glucose. **C.** Average AUC of the 2-hr OGTT excursion profiles for male mice at 4, 5, 6, 7, and 14 weeks of age. Oral glucose tolerance test (OGTT) performed in 4 (**D**) and 14-week-old (**E**) female mice following a 4-hour fast in response to oral gavage of 2g/kg glucose. **F.** Average AUC of the 2-hr OGTT excursion profiles for female mice at 4, 5, 6, 7, and 14 weeks of age. Data are presented as mean±SEM; N=5 mice/cohort. Data were analyzed using student's t-test; *P<0.05.

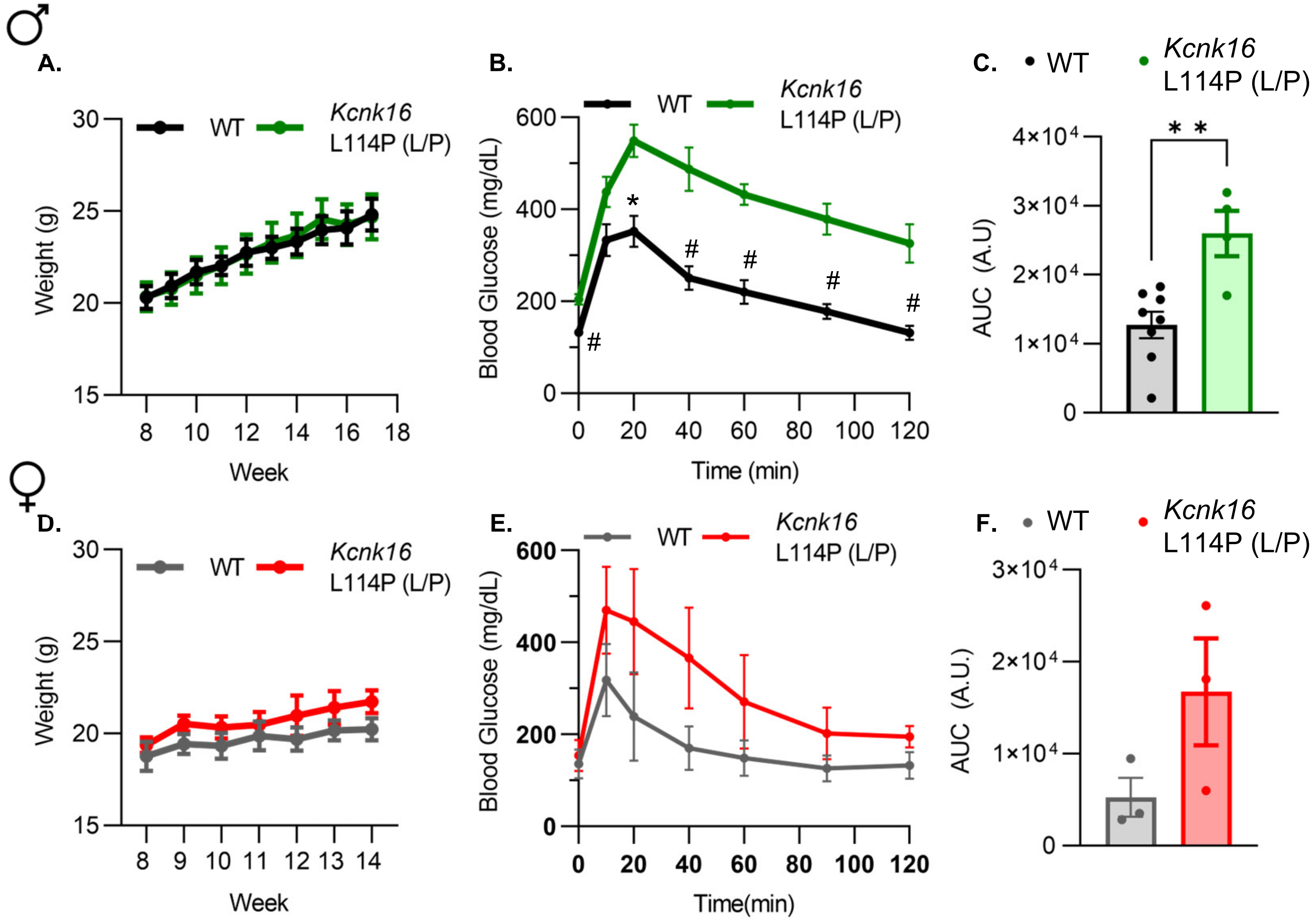
♂
A.
B.
C.
WT
Kcnk16
L114P (L/P)
Weight (g)
Week
Blood Glucose (mg/dL)
Time (min)
AUC (A.U)
**
♀
D.
E.
F.
Time(min)
AUC (A.U.)

**Supplemental figure 4. Glucose homeostasis is impaired in *Kcnk16* L114P (L/P) mice in the B6 background. A.** Body weight measurements of male WT (black; N=8) and *Kcnk16* L114P (L/P; green; N=4) mice in the C57Bl/6J background. **B.** Intraperitoneal glucose tolerance test (i.p. GTT) performed in 10-week-old male mice following a 4-hour fast in response to 2mg/g glucose injection. **C.** Average AUC of the 2-hr GTT excursion profiles in (**c**). **D.** Body weight measurements of female WT (gray; N=3) and *Kcnk16* L114P (L/P; red; N=3) mice. **E.** I.P. GTT performed in 11-week-old female mice following a 4-hour fast in response to 2mg/g glucose injection. **F.** Average AUC of the 2-hr GTT excursion profiles in (**E**). Data are presented as mean±SEM. Data were analyzed using student's t-test; **P<0.01, #P<0.001.

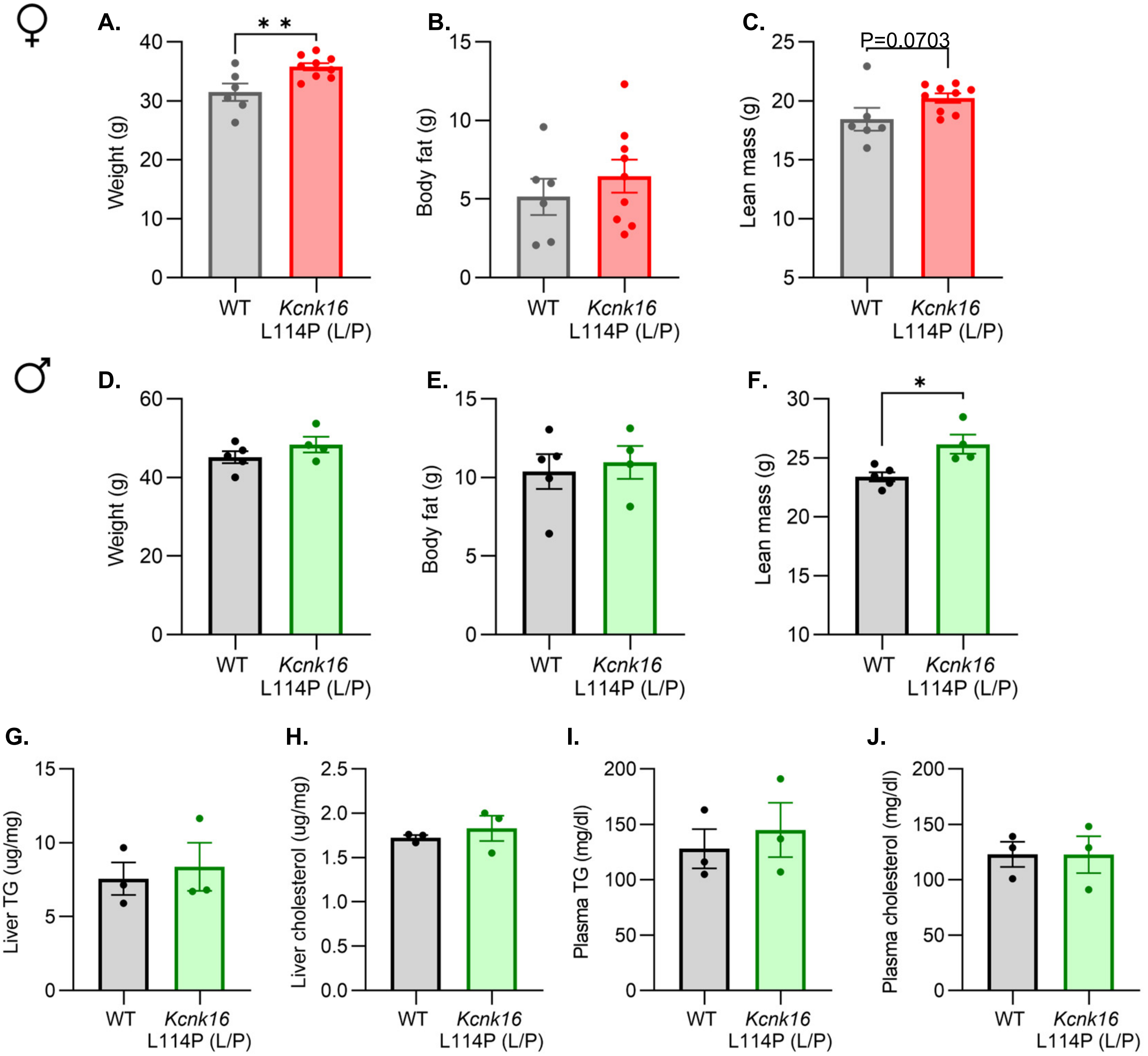

♀
A.
* *
Weight (g)
40
30
20
10
0
WT
Kcnk16
L114P (L/P)
B.
Body fat (g)
15
10
5
0
WT
Kcnk16
L114P (L/P)
C.
P=0.0703
Lean mass (g)
25
20
15
10
5
WT
Kcnk16
L114P (L/P)
♂
D.
Weight (g)
60
40
20
0
WT
Kcnk16
L114P (L/P)
E.
Body fat (g)
15
10
5
0
WT
Kcnk16
L114P (L/P)
F.
*
Lean mass (g)
30
25
20
15
10
WT
Kcnk16
L114P (L/P)
G.
Liver TG (ug/mg)
15
10
5
0
WT
Kcnk16
L114P (L/P)
H.
Liver cholesterol (ug/mg)
2.5
2.0
1.5
1.0
0.5
0.0
WT
Kcnk16
L114P (L/P)
I.
Plasma TG (mg/dl)
200
150
100
50
0
WT
Kcnk16
L114P (L/P)
J.
Plasma cholesterol (mg/dl)
200
150
100
50
0
WT
Kcnk16
L114P (L/P)

**Supplemental figure 5. Body composition measurements and assessment of plasma and liver triglycerides and total cholesterol. A.-F.** Body composition analysis of male and female B6;CD-1 WT and *Kcnk16* L114P (L/P) mice assessing weight (g), body fat (g), and lean mass (g) (N=5-9 mice/genotype). **G.-J.** Average liver and plasma cholesterol and triglyceride levels in male B6;CD-1 WT and *Kcnk16* L114P (L/P) mice (N=3/genotype). Data are presented as mean±SEM. Data were analyzed using student's t-test.

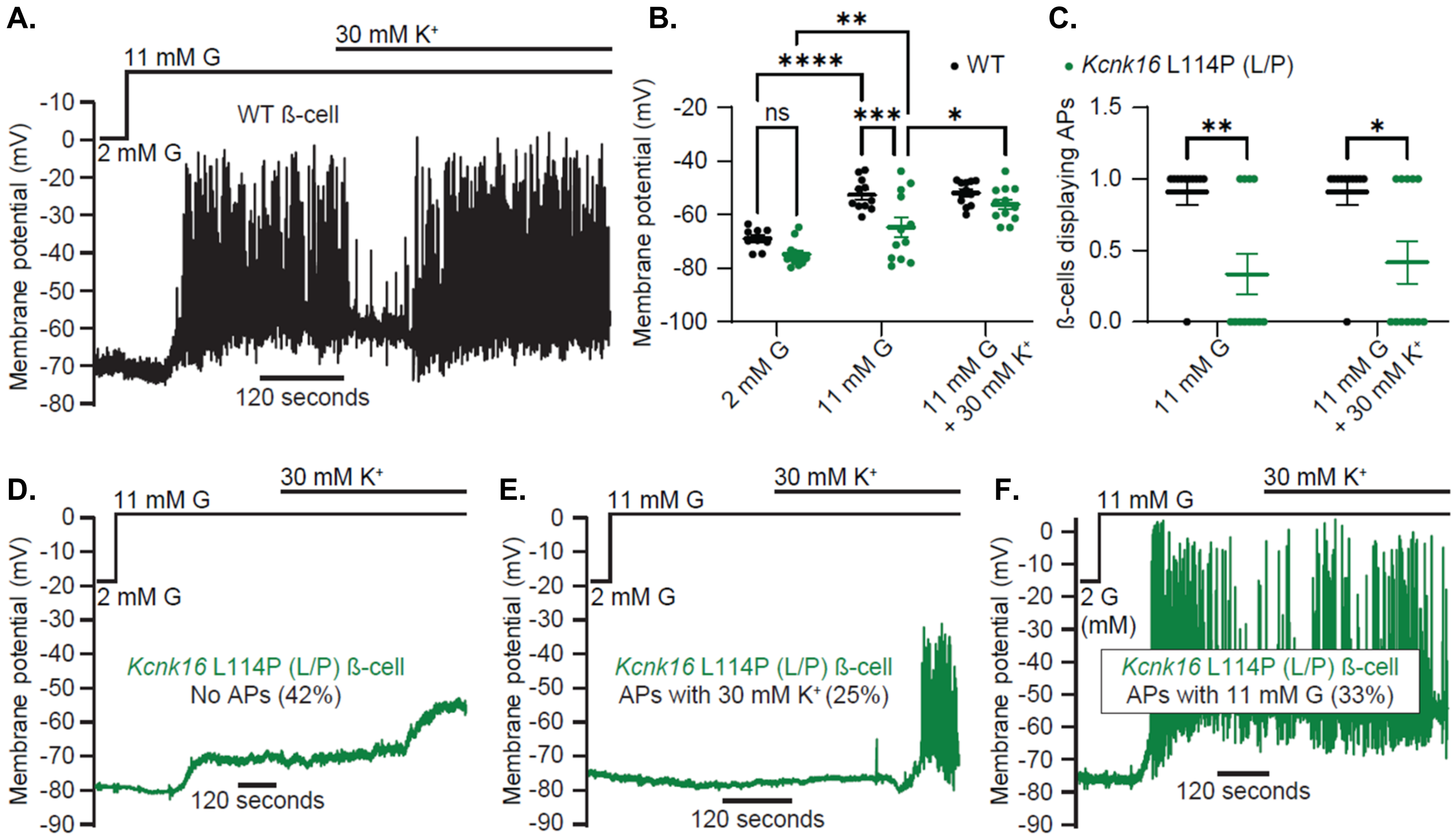
A.
30 mM K+
11 mM G
WT ß-cell
2 mM G
Membrane potential (mV)
120 seconds
B.
• WT
• Kcnk16 L114P (L/P)
ns
****
**
***
*
Membrane potential (mV)
2 mM G
11 mM G
11 mM G + 30 mM K+
C.
ß-cells displaying APs
**
*
11 mM G
11 mM G + 30 mM K+
D.
11 mM G
30 mM K+
2 mM G
Kcnk16 L114P (L/P) ß-cell
No APs (42%)
120 seconds
E.
Kcnk16 L114P (L/P) ß-cell
APs with 30 mM K+ (25%)
120 seconds
F.
2 G (mM)
Kcnk16 L114P (L/P) ß-cell
APs with 11 mM G (33%)
120 seconds

**Supplemental Figure 6. *Kcnk16* L114P attenuates β-cell glucose-stimulated electrical excitability in a heterogenous manner. A.** Representative perforated patch-clamp $V_m$ recording in response to 11 mM G and 11 mM G+30 mM $K^+$ from a WT ß-cell expressing GCaMP6s driven by an optimized RIP within an islet cluster (representative of N=11 islets from 5 mice). **B.** Average resting $V_m$ of GCaMP6s positive WT and *Kcnk16* L114P (L/P) ß-cells within islet clusters in the presence of 2 mM G as well as plateau $V_m$ in the presence of 11 mM G and 11 mM G+30 mM $K^+$. **C.** Percentage of GCaMP6s positive WT and *Kcnk16* L114P (L/P) ß-cells displaying action potentials (APs) in response to 11 mM G and 11 mM G+30 mM $K^+$. **D.** Representative perforated patch-clamp $V_m$ recordings in response to 11 mM G and 30 mM $K^+$ illustrating cell-to-cell heterogeneity from *Kcnk16* L114P (L/P) ß-cells expressing GCaMP6s driven by an optimized RIP (representative of N=12 islets from 5 mice). Data are presented as mean±SEM. Data were analyzed using two-way ANOVA; *P<0.05, **P<0.01, ***P<0.001, and ****P<0.0001.

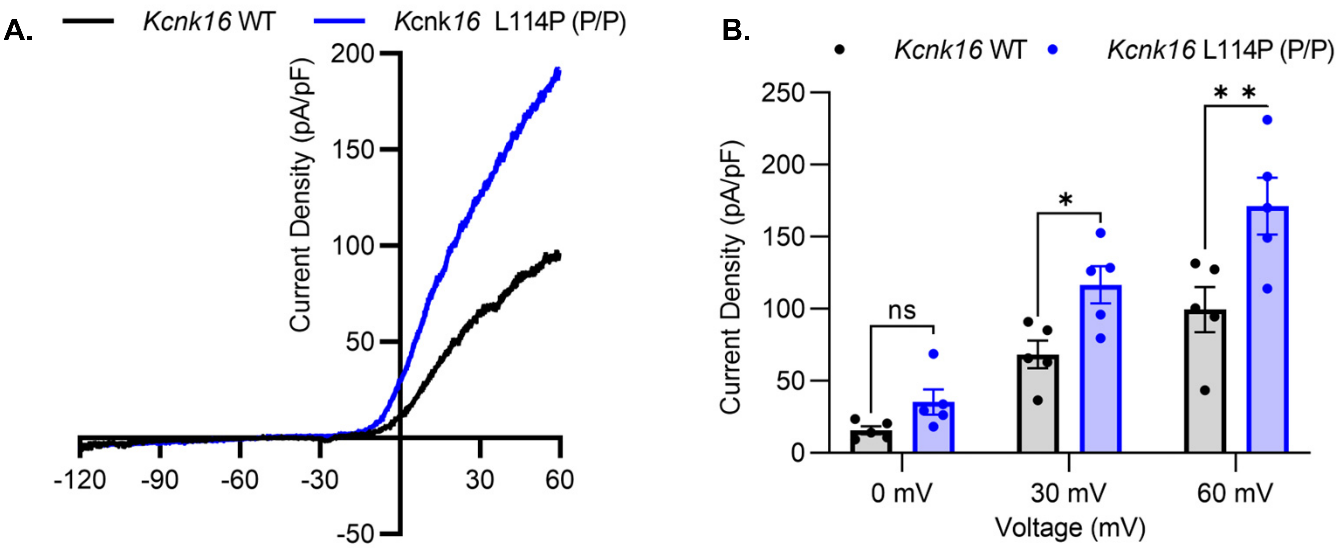
A.
Kcnk16 WT
Kcnk16 L114P (P/P)
Current Density (pA/pF)
200
150
100
50
-50
-120
-90
-60
-30
30
60
B.
Kcnk16 WT
Kcnk16 L114P (P/P)
Current Density (pA/pF)
250
200
150
100
50
0
ns
*
* *
0 mV
30 mV
60 mV
Voltage (mV)

**Supplemental figure 7. Two-pore domain $K^+$ channel currents in β-cells from homozygous *Kcnk16* L114P (P/P) mice also exhibit a modest increase. A.** Representative whole-cell two-pore domain $K^+$ channel current density (pA/pF) recorded using a voltage ramp (-120 mV to +60 mV) in 11 mM G in islets from B6; CD-1 WT and *Kcnk16* L114P (P/P) P4 neonates. B. Average current density (pA/pF) measured at the specified membrane potentials (N=5 cells/genotype). Data are presented as mean±SEM. Data were analyzed using two-way ANOVA.

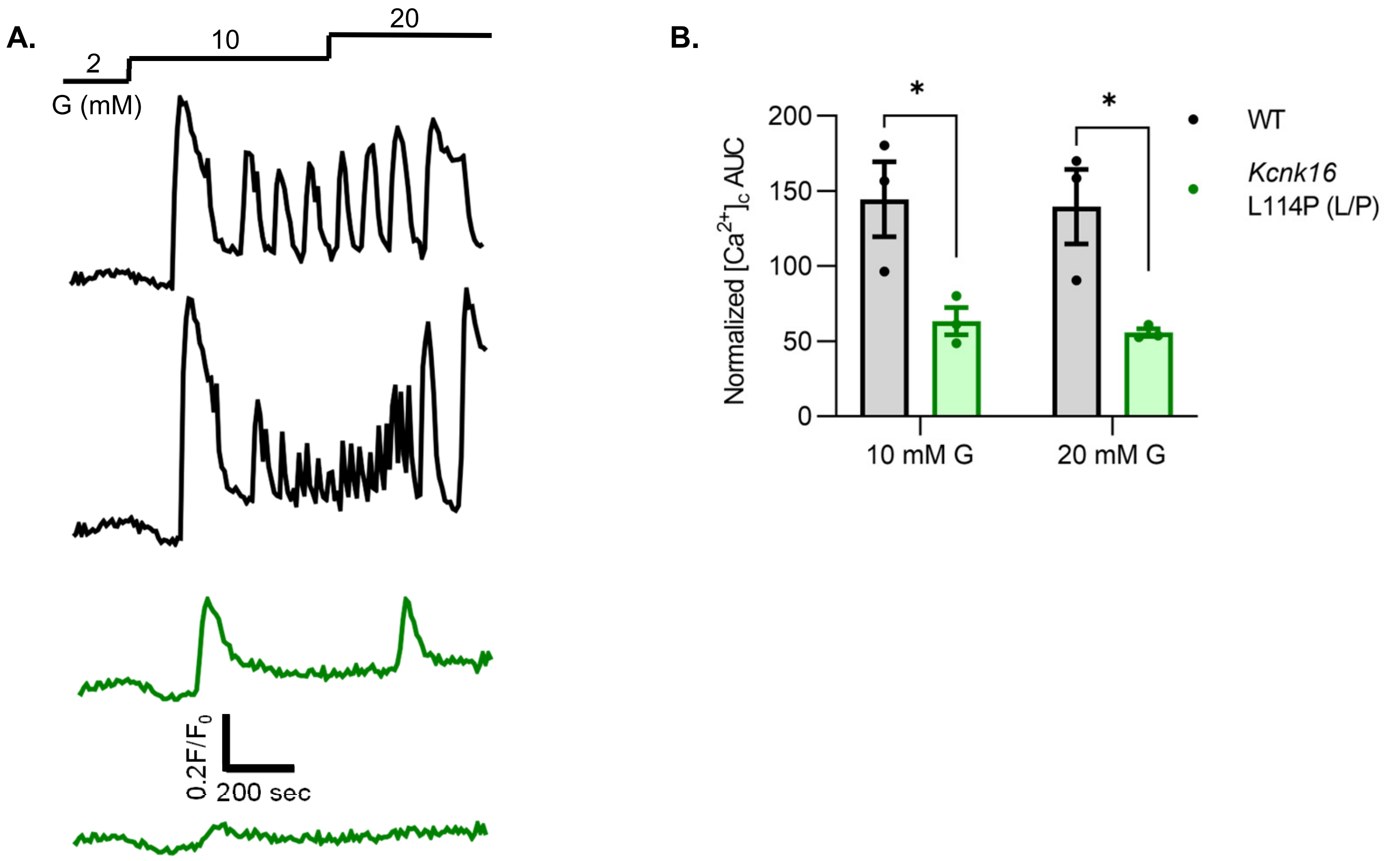

A.
2
10
20
G (mM)
0.2F/F0
200 sec
B.
*
*
200
150
100
50
0
Normalized [Ca2+]c AUC
10 mM G
20 mM G
WT
Kcnk16
L114P (L/P)

**Supplemental figure 8. Islets from *Kcnk16* L114P (L/P) mice on the B6 background also exhibit blunted glucose-stimulated $Ca^{2+}$ entry. A.** Representative glucose-stimulated $[Ca^{2+}]_c$ influx traces in islets from male WT and *Kcnk16* L114P(L/P) mice in the C57BL/6J background in response to 2 mM G, 10 mM G, and 20 mM G (N=3 mice/genotype). **B.** Average total AUC in response to the indicated glucose concentrations in islets from male WT and *Kcnk16* L114P(L/P) mice. Data are presented as mean±SEM. Data are analyzed using student's t-test.

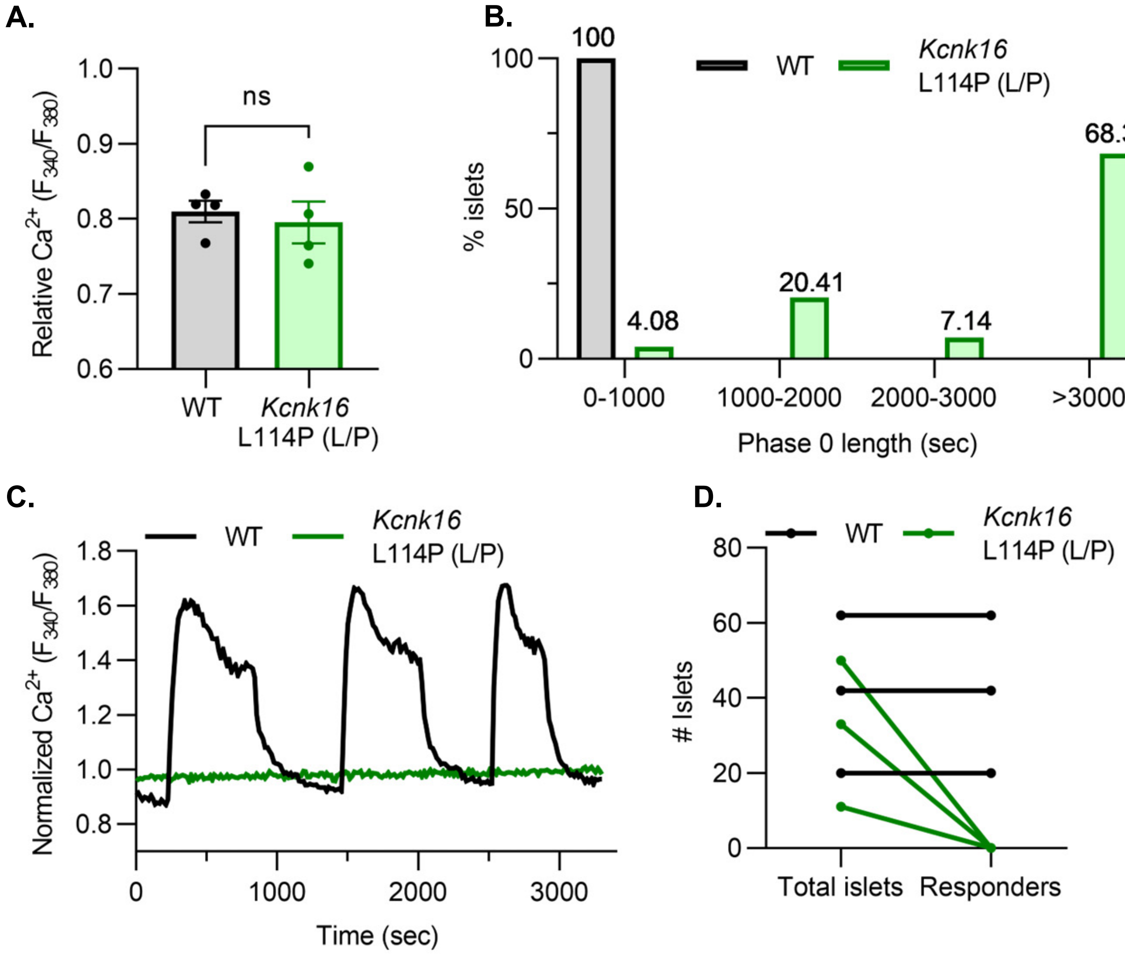
A.
ns
Relative Ca2+ (F340/F380)
1.0
0.9
0.8
0.7
0.6
WT
Kcnk16
L114P (L/P)
B.
100
WT
Kcnk16
L114P (L/P)
68.37
4.08
20.41
7.14
% islets
100
50
0
0-1000
1000-2000
2000-3000
>3000
Phase 0 length (sec)
C.
WT
Kcnk16
L114P (L/P)
Normalized Ca2+ (F340/F380)
1.8
1.6
1.4
1.2
1.0
0.8
0
1000
2000
3000
Time (sec)
D.
WT
Kcnk16
L114P (L/P)
### Islets
80
60
40
20
0
Total islets
Responders

**Supplemental figure 9. *Kcnk16* L114P (L/P) islets exhibit prolonged glucose-stimulated phase 0 $[Ca^{2+}]_{ER}$ uptake and show a complete absence of $Ca^{2+}$ oscillations. A.** Average relative $[Ca^{2+}]_c$ at 2 mM G in islets from B6; CD-1 WT and *Kcnk16* L114P (L/P) mice (N=4 mice/genotype). **B.** Percent islets that exhibit the corresponding phase 0 response (drop in $[Ca^{2+}]_c$) length (sec) in WT and *Kcnk16* L114P (L/P) mice. **C.** Representative glucose-stimulated $[Ca^{2+}]_c$ oscillations recorded at 9 mM G in islets from WT and *Kcnk16* L114P (L/P) mice. **D.** Total number of islets analyzed for $[Ca^{2+}]_c$ oscillations vs. the number of islets that exhibited $[Ca^{2+}]_c$ oscillations from WT and *Kcnk16* L114P (L/P) mice (N=3 mice/genotype). Data are presented as mean±SEM. Data are analyzed using student's t-test.

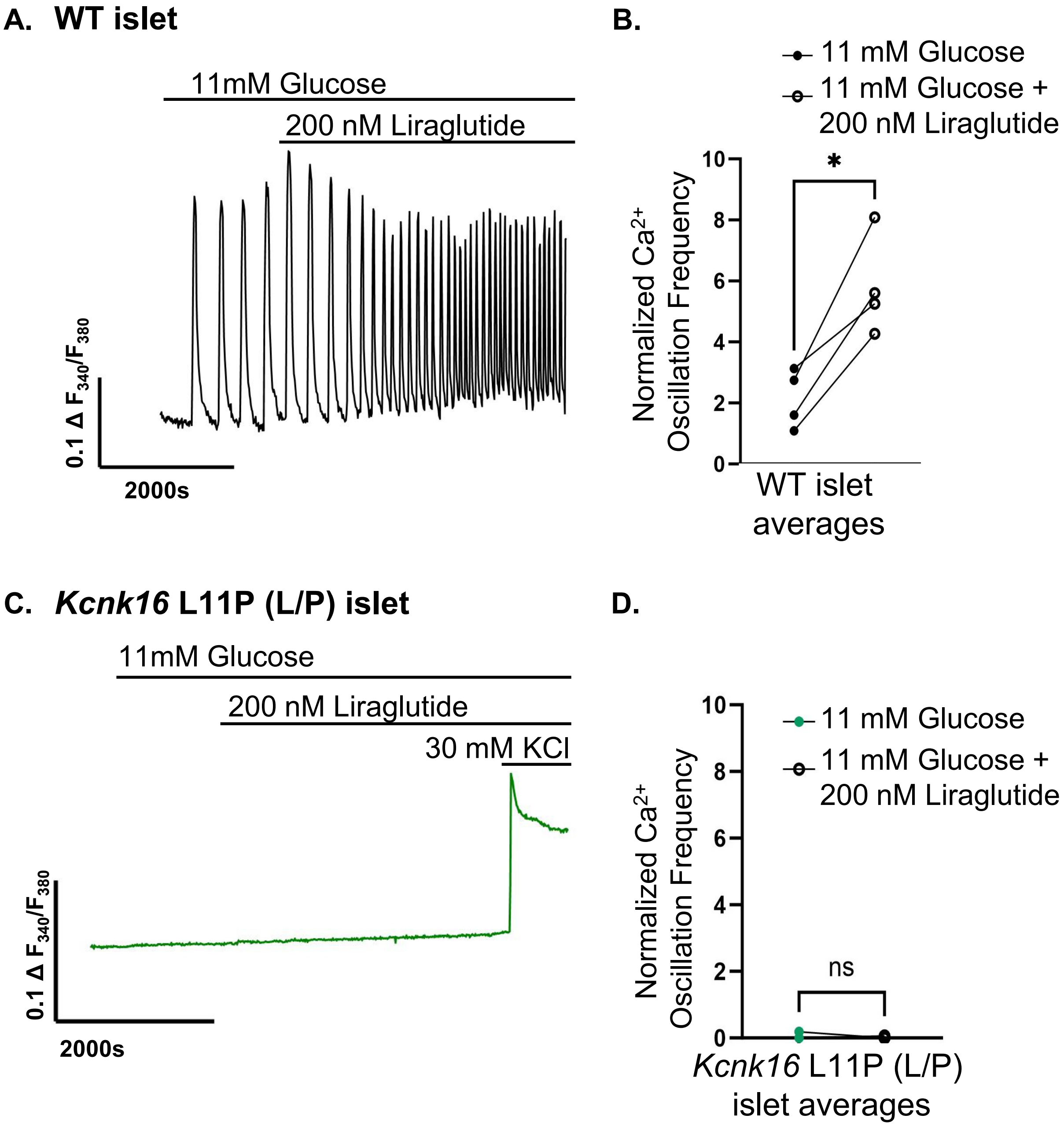

A. WT islet
11mM Glucose
200 nM Liraglutide
0.1 Δ F340/F380
2000s
B.
11 mM Glucose
11 mM Glucose + 200 nM Liraglutide
Normalized Ca2+ Oscillation Frequency
*
WT islet averages
C. Kcnk16 L11P (L/P) islet
11mM Glucose
200 nM Liraglutide
30 mM KCl
0.1 Δ F340/F380
2000s
D.
11 mM Glucose
11 mM Glucose + 200 nM Liraglutide
Normalized Ca2+ Oscillation Frequency
ns
Kcnk16 L11P (L/P) islet averages

**Supplemental figure 10. Liraglutide increased $Ca^{2+}$ oscillation frequency in WT islets but does not impact *Kcnk16* L114P (L/P) mouse islet $Ca^{2+}$ handling. A.** Representative Islet $Ca^{2+}$ recording from a WT islet in 11 mM glucose treated with 200 nM liraglutide. **B.** Average Ca2+ oscillation frequency of WT islets in 11 mM glucose and in 11 mM glucose with 200 nM Liraglutide (LIRA, N = 57 islets from 4 mice). **C.** Representative Islet $Ca^{2+}$ recording from a *Kcnk16* L114P mice islet in 11 mM glucose treated first with 200 nM liraglutide and then with 30mM KCl. **D.** Average $Ca^{2+}$ oscillation frequency of *Kcnk16* L114P islets in 11 mM glucose and in 11 mM glucose with 200 nM Liraglutide (LIRA, N = 80 islets from three mice). Data are presented as mean±SEM. Data were analyzed using paired Student's t-test.

A.

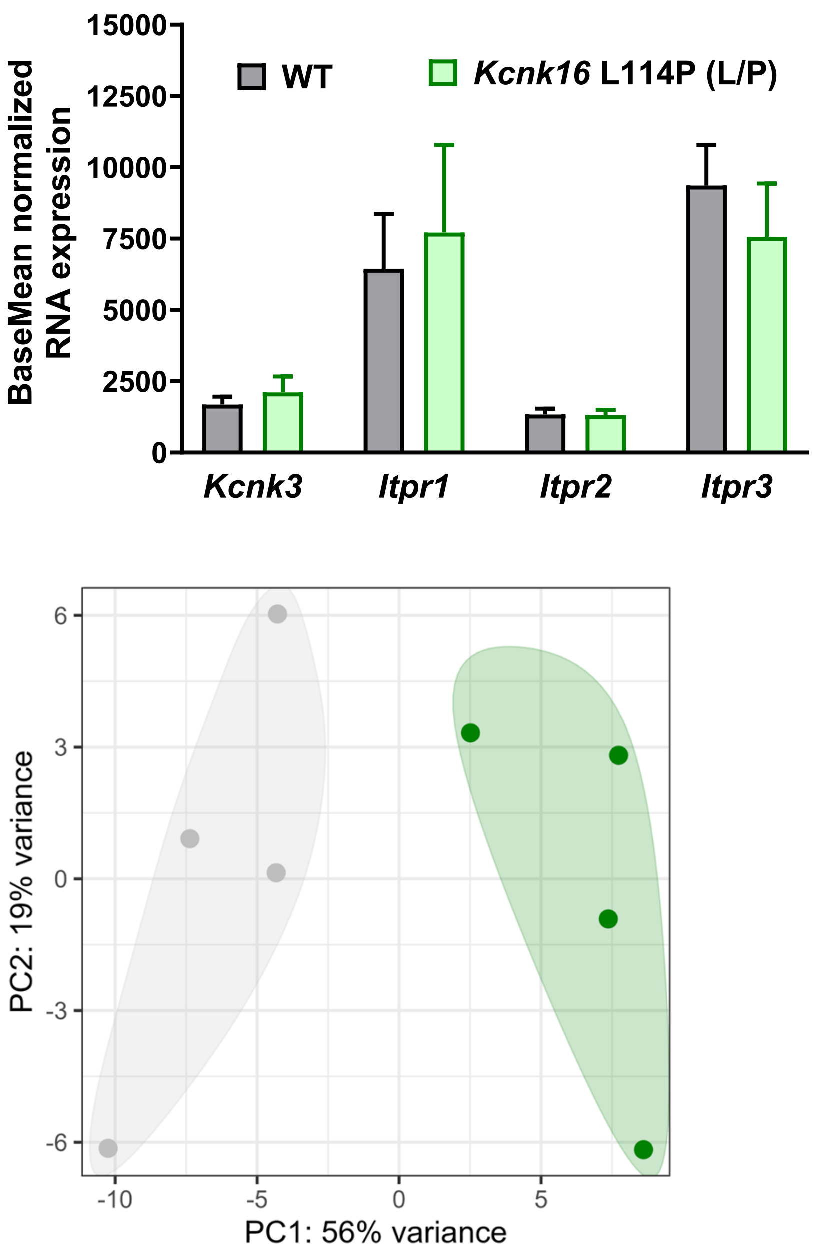

BaseMean normalized RNA expression
15000
12500
10000
7500
5000
2500
0
WT
*Kcnk16* L114P (L/P)
*Kcnk3*
*Itpr1*
*Itpr2*
*Itpr3*
PC2: 19% variance
6
3
0
-3
-6
-10
-5
0
5
PC1: 56% variance


B.

**Supplemental figure 11. RNA sequencing showed equivalent expression of a subset of genes encoding proteins known to control $[Ca^{2+}]_{ER}$ in WT and *Kcnk16* L114P (L/P) male islets**. **A.** Average normalized RNA levels (baseMean) of *Kcnk3*, *Itpr1*, *Itpr2*, and *Itpr3* from WT islets (grey bars, N=4) and *Kcnk16* L114P (L/P) (green bars, N=4). Data are presented as mean±SEM. **B.** Principal component analysis (PCA) showing clustering of WT (gray) and *Kcnk16* L114P (L/P; green) islet RNA samples.

| Gene | Forward primer | Reverse primer |
|---|---|---|
| *18sRNA* | GTAACCCGTTGAACCCCATT | CCATCCAATCGGTAGTAGCG |
| *Cacna1g* | GAGACACAGAGTACGGGAGC | CAGGCATTTCATGGTCAGCG |
| *Sst* | CCACCGGGAAACAGGAACTG | TTGCTGGGTTCGAGTTGGC |
| *Asb11* | TGGTGGACTGTCAGACTGCT | ATTGACGTTGATGCCTTGCG |
| *Fxyd3* | ACTCTGCTTTCTCCCGGAAC | CTCGGAGGCTGTACCAATCATA |
| *Aldh1a3* | GGGTCACACTGGAGCTAGGA | CTGGCCTCTTCTTGGCGAA |
| *Camk1d* | CCGCCCTACAGCATTAGTCT | GAAAAGGCCCCAGTTCCGA |
| *Cxcl1* | ACCCAAACCGAAGTCATAGCC | TTGTCAGAAGCCAGCGTTCA |
| *Adcyap1r1* | CTGCGTGCAGAAATGCTACTG | AGCCGTAGAGTAATGGTGGATAG |
| *Aldob* | AGAAGGACAGCCAGGGAAAT | GTTCAGAGAGGCCATCAAGC |
| *Pdk4* | TGGTAGCAGTAGTCCAAGATGC | GTGGATTGGTTGGCCTGGAA |
| *Tgfb2* | TCGACATGGATCAGTTTATGCG | CCCTGGTACTGTTGTAGATGGA |

**Supplemental table 1. Mouse primer sequences used for qRT-PCR.**
